# Force and the *α*-C-terminal domains bias RNA polymerase recycling

**DOI:** 10.1101/2023.11.02.565415

**Authors:** Jin Qian, Bing Wang, Irina Artsimovitch, David Dunlap, Laura Finzi

**Affiliations:** Physics Department, Emory University, GA, USA; Department of Microbiology, The Ohio State University, OH, USA

**Keywords:** transcription, magnetic tweezers, RNA polymerase, force spectroscopy

## Abstract

After RNA polymerase (RNAP) reaches a terminator, instead of dissociating from the template, it may diffuse along the DNA before restarting RNA synthesis from the previous or a different promoter. We monitored such secondary transcription using magnetic tweezers to determine the effect of low forces, protein roadblocks, and transcription factors. Up to 60% of RNAPs diffused along the DNA after termination. Force biased the direction of diffusion (sliding) and the velocity increased rapidly with force up to 0.7 pN and much more slowly thereafter. Sigma factor 70 (***σ*^70^**) likely remained bound to these sliding RNAPs, enabling recognition of secondary promoters and additional rounds of transcription, and the addition of elongation factor NusG, which competes with ***σ*^70^** for binding to RNAP, limited additional rounds of transcription. Surprisingly, RNAP blocked by a DNA-bound *lac* repressor could slowly re-initiate transcription at the blockage and was not affected by NusG, suggesting a ***σ***-independent pathway. Force biased the frequency of encounters between convergent or divergent promoters such that transcription was repetitive only from the promoter opposing the direction of force. Sliding RNAP recognized and could subsequently re-initiate from promoters in either orientation. However, deletions of the ***α***-C-terminal domains severely limited the ability of RNAP to turn around.

## 1 Introduction

A transcription cycle comprises recognition of the promoter sequence by RNAP, initiation, elongation, and termination. In canonical termination, RNAP dissociates from the DNA template at a terminator and diffuses away, predominantly in 3-D [1, 2]. However, previous reports indicate that upon reaching a terminator RNAP may remain on the DNA template [3, 4] and diffuse one-dimensionally to a nearby promoter to execute another cycle of transcription in the same or opposite direction [5–7]. The ability of RNAP to repetitively transcribe the same DNA sequence by sliding back to the promoter might contribute significantly to gene regulation. Indeed, cycles of transcription by the same RNAP enzyme would efficiently accumulate transcript and eliminate the need to repeatedly recruit RNAP and reduce the probability of collisions among different RNAPs.

While this alternative to canonical termination has been reported in different contexts [4–7], the biological significance and mechanism of repetitive transcription remains poorly understood. Force could influence this process by biasing the direction of RNAP diffusion, which might be blocked by DNA-bound proteins as well. The role of sigma factor (*σ*) in repetitive transcription is also unclear. In order to initiate transcription, RNAP requires *σ* factor for open complex formation [8]. While the *σ* factor is thought to detach from RNAP after initiation, there is evidence that *σ* may remain associated with transcription elongation complexes and promote open bubble formation during secondary transcription events [9]. Moreover, it is also unclear how an RNAP, one-dimensionally sliding along DNA after completing a “primary” round of elongation in one direction, might switch orientation to initiate “secondary” transcription at a promoter oriented in the opposite direction.

In this study, *in vitro* assays using single-molecule magnetic tweezers (MTs) were used to observe transcribing RNAPs following arrival at terminators in different buffer conditions with various forces on the enzyme. Under various experimental conditions 20% to 60% of RNAPs exhibited non-canonical termination, but multiplexed MT assays produced statistically significant data for analyses of the effect of force, roadblock proteins, and different terminators on repetitive transcription. External force directed and modified the velocity of diffusion, biasing the search for a secondary promoter. The *σ* factor was likely required for relatively rapid initiation of repetitive transcription from promoters but not for slower re-initiation from a non-promoter site. While sliding RNAP re-initiated from promoters oriented in either direction with respect to previous elongation, deletion of the C-terminal domains of the *α* subunits (*α*-CTD), known to interact with DNA, nearly abolished the switch of direction.

## 2 Results

### 2.1 Force directs RNAP diffusion and repetitive transcription

To investigate the mechanism underlying repetitive transcription by RNAP, we applied mechanical force to hinder or enhance RNAP translocation on a DNA template containing a promoter, a high affinity *lac* repressor binding site (Lac O1), and a terminator sequence. MTs were used to simultaneously monitor multiple DNA templates with single transcription complexes. The orientation of the promoter sequence(s) determined whether external force opposed or assisted transcription, and the magnitude of force was dependent on the separation between the magnet and the paramagnetic particle linked to RNAP, as illustrated in Fig. 1a. To examine the effect of force, the experiments were conducted at forces ranging from −5 (opposing) to +5 (assisting) pN. After reaching a terminator, opposing force drove single, diffusing RNAPs to slide back to the promoter where some enzymes re-initiated transcription (Fig. 1b, cycles 1 & 2). Single RNAPs repetitively transcribed the same sequence as many as seven times. Finally, a recycling RNAP slid to the promoter where it paused briefly before either dissociating directly from the promoter or sliding off the end of the DNA template (Fig. 1b, cycle 3). These two pathways of RNAP release could not be distinguished in the assay, because the bead rapidly separated from the surface in either case.

**Figure 1.**
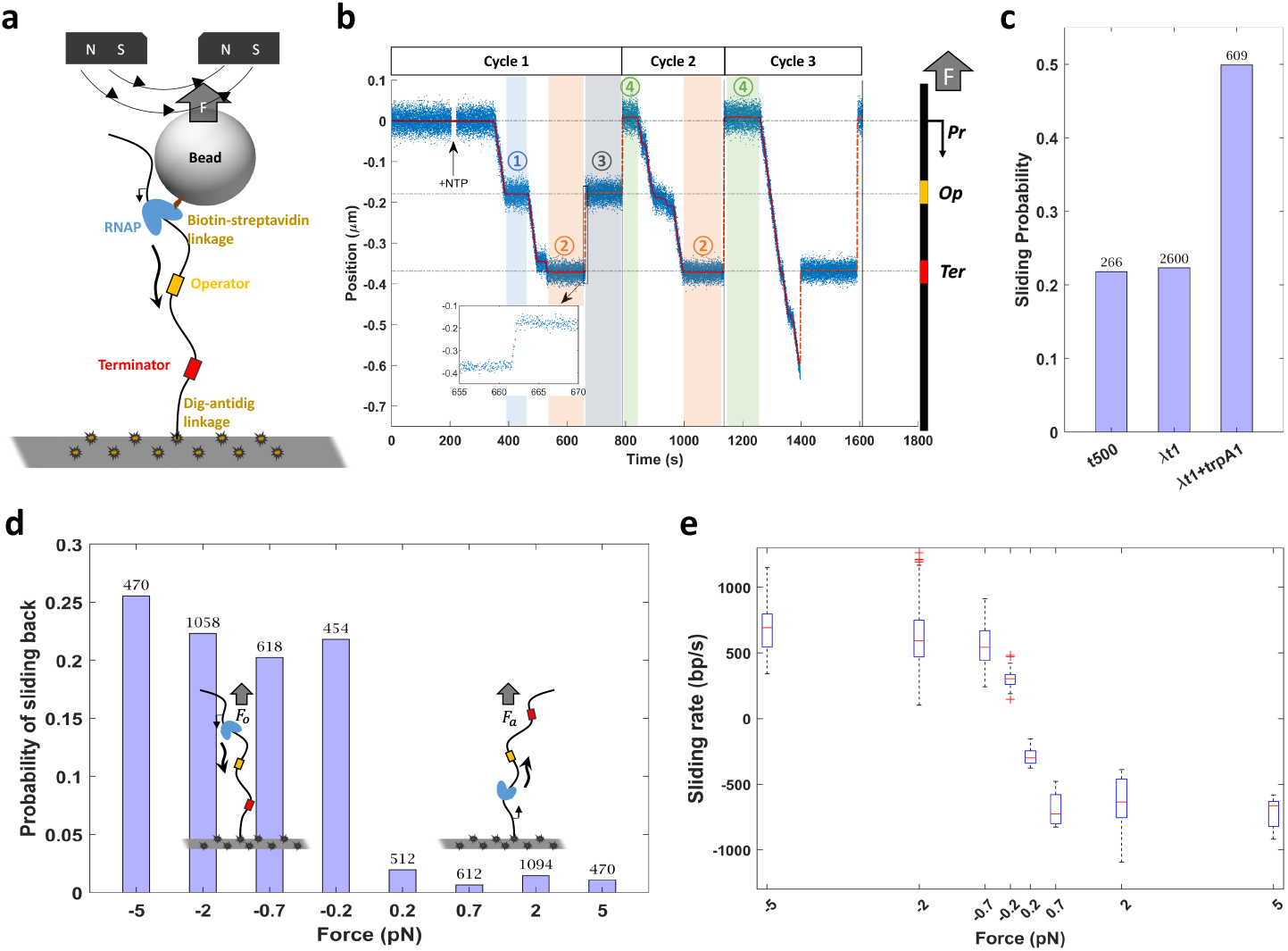
Force-directed sliding leads to repetitive transcription. (a) A diagram of the experimental setup for transcription against opposing force. (b) A representative recording of multiple rounds of transcription under opposing force includes a temporary roadblock-associated pause during transcription (shaded region 1), pauses at the terminator (shaded regions 2), RNAP temporarily roadblocked during backward sliding (shaded region 3), and pauses at the promoter prior to re-initiation (shaded regions 4). The inset shows data points corresponding to RNAP sliding back from the terminator in cycle 1. (c) On templates with a dual terminator sequence, the percentage of RNAP that slid backward was twice that on templates with single terminators. The total number of events are listed above each bar. (d) Opposing force (negative values) significantly raised the probability that the post-terminator complex slid toward the promoter from which the previous cycle of transcription initiated. The total number of events are listed above each bar. (e) RNAP sliding rates increased rapidly as opposing (-) or assisting (+) force increased from 0 to 0.7 pN but plateaued thereafter. Red crosses indicate outliers.

Repetitive transcription is an intriguing alternative to canonical termination in which RNAP dissociates from the DNA template. Under opposing force, slightly more than 20% of RNAPs slid backward from single *λ*T1 or T500 terminators. However, a pair of consecutive terminators (Fig. 1c) doubled the probability of sliding to *∼*50%. This suggests that an encounter with a terminator favors the transition from a transcribing to a sliding conformation. Although sliding most often began from a terminator, occasionally it started from a position beyond the terminator (Fig. 1b Cycle 3). In these cases, conformational changes associated with termination that promote sliding might have occurred slowly during run-on elongation. This is consistent with a reported two-stage post-elongation complex (post-termination complex) [10].

Repetitive transcription was sensitive to the direction but not the magnitude of force. Under opposing forces up to 25% of post-termination complexes rapidly slid backwards to the previously utilized promoter (Fig. 1d) and *∼*10% of these re-initiated transcription (Table S1 single promoter records summary). Even under forces as low as 0.2 pN, post-termination complexes almost always slid in the direction of applied force. To further test the effect of force on sliding, a reversed DNA construct was employed (Fig. 1d right). After transcribing under assisting force, post-termination complexes rarely slid backward against the applied force, irregardless of the magnitude. This susceptibility to force indicates that post-termination complexes were driven to the promoter by 1D-diffusion along the DNA template instead of 3D diffusion or inter-segment transfer. Furthermore, while the rates of transcription were consistent with literature data [11–13], the distributions of rates measured in successive cycles associated with one template were almost identical and distinct from those associated with another identical template (Fig. S1e) as reported previously [13]. Together these facts strongly indicate that the same RNAP enzyme produced the repetitive cycles of transcription in any given recording.

Unidirectional sliding of RNAP would be expected if force were to bias one-dimensional diffusion. While bidirectional diffusion reported previously persisted for minutes [5, 7], fast, unidirectional sliding toward a promoter required only *<*3 sec even under extremely low force, 0.2 pN. The magnitude of applied force did not change the probability of sliding, but higher force increased the sliding velocity (Fig. 1e). It was estimated to be at least 200 bp/sec at the weakest forces applied and increased with force magnitude. The sliding velocity increased linearly with the magnitude of low forces (*<*2pN) but plateaued at higher forces. The conformational entropy of the free end of the DNA template feeding into a sliding RNAP might be one factor limiting the sliding velocity. For DNA threading into a 5 nm long, 10 nm diameter pore, the associated entropic force of uncoiling for threading has been estimated to be *∼*0.7-1.9 pN [14], which coincides well with the inflection point in the sliding velocity versus force data. Forces opposing or assisting transcription clearly bias the direction of post-termination complex sliding and reduce delays between successive encounters with nearby promoters.

### 2.2 Secondary transcription from a roadblock site

When LacI protein was introduced, RNAP paused frequently at the O1 binding site during transcription, confirming that LacI acts as a “roadblock” (Fig. 2a Cycle 1). LacI also blocked post-termination sliding (Fig. 2a Cycle 2, Fig. S2a), and the distribution of dwell times associated with roadblock-induced pauses did not depend on the magnitude of the force (Fig. S2b). This result further supports the idea that force-driven post-termination complexes slide along the DNA template, pausing until roadblocks dissociate from the DNA.

**Figure 2.**
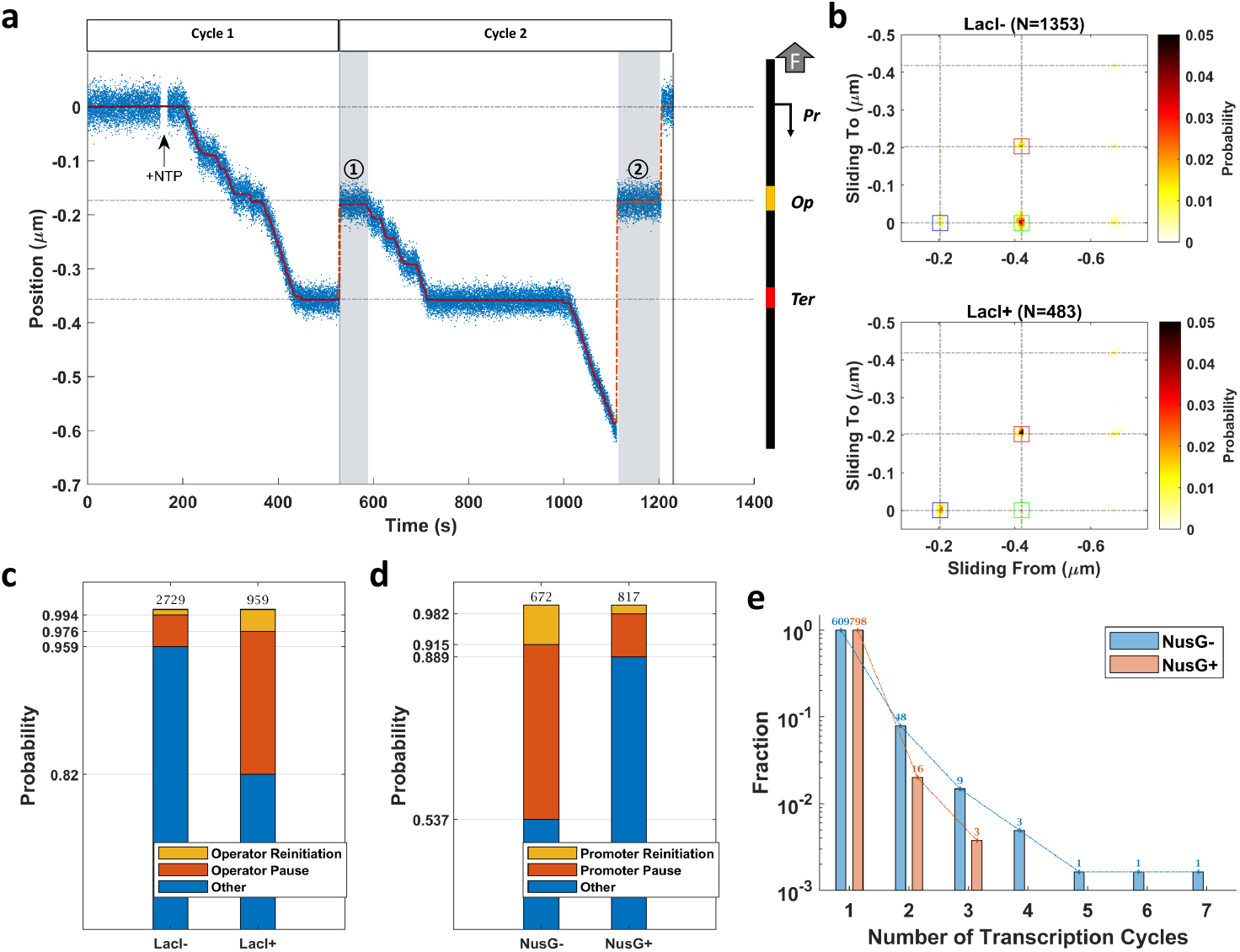
DNA-binding protein roadblocks and NusG affected post-termination sliding and repetitive transcription. (a) In this representative recording, the shaded regions indicate pauses at the roadblock during post-termination sliding followed by re-initiation (1) or continued sliding past the roadblock to the promoter (recapture) (2). (b) Heat maps indicate probabilities associated with locations at which post-termination complexes started and stopped sliding under opposing force in buffer without (upper) or with (lower) LacI. Sliding primarily began at the terminator (*∼*-0.4 µm) and ended at promoter (*∼*0 µm) (green box), unless LacI was present to block sliding and induce terminator-to-LacI binding site (red box) and LacI binding site-to-promoter (blue box) sliding events. (c) Adding LacI to the buffer increased the probability of roadblocking and re-initiation at the road-block. (d) Although NusG diminished the probability of sliding to promoters, it did not change the ratio of re-initiation, *∼*0.1 (yellow/red). (e) A minor population of RNAPs, less than 10%, exhibited multiple cycles of transcription.

After pauses at the roadblock, most post-termination complexes slid past to the promoter where *∼*4% recommenced transcription independently of whether LacI was present (Fig. S3a). Unexpectedly, when LacI was present, *∼*2.4% of sliding post-termination complexes recommenced transcription at the roadblock while significantly less, *∼*0.6%, did so in the absence of LacI (Fig. 2c). Remarkably, the dwell times of post-termination complexes that re-initiated transcription at the roadblock were much longer than those that paused and then continued sliding (Fig. S2b). Evidently, after sufficient time at roadblocks, post-termination complexes may form open complexes and recommence transcription. Thus, protein roadblocks on DNA templates can block elongation complexes to delay transcript completion or produce 3’-truncations [15] as well as post-termination complexes to produce 5’-truncated transcripts.

### 2.3 Sigma factor promoted sliding and secondary promoter recognition

Promoter recognition by RNAP and transition to an open complex involves *σ* factors. In a buffer without free *σ*^70^, secondary initiation from the promoter would require *σ* factor to remain bound to RNAP throughout primary transcription and subsequent post-termination complex sliding to the promoter. To test this hypothesis, NusG, a transcription elongation factor that competes with *σ*^70^ for binding to RNAP [16], was introduced. NusG diminished the frequency of repetitive transcription events from the promoter site (Fig. 2d) as expected for competition that would release *σ*^70^ from elongation and post-termination complexes. More precisely, NusG decreased the probability of post-termination complexes sliding rapidly along DNA template, but a similar fraction of sliding post-termination complexes re-initiated after reaching the promoter (Fig. 2d). This is consistent with reports that *σ*^70^ often remains associated with post-termination complexes on DNA [7, 9], and another report describing experiments in which optical tweezers were used to apply forces to core RNAP and found only brief, force-sensitive dwell times at t500, *his*, and *λ*T2 terminators [17]. In the current experiments, the probability of re-initiation dropped precipitously after one round (Fig. 2e). With a measured dissociation constant of 3.8*±*0.8 *×* 10*^−^*^3^ [9], few RNAPs were likely to retain *σ*^70^ through multiple cycles of transcription lasting several hundred seconds (Fig. 1b).

Similarly, NusG negligibly changed re-initiation at the roadblock (Fig. S3a), although post-termination complexes lingered much longer at roadblocks than at promoters before re-initiating transcription (Fig. S2b). This suggests that core RNAP enzyme, without a *σ* factor, can form a transcription bubble for elongation at a non-promoter sequence, albeit more slowly.

### 2.4 Force biased re-initiation between converging or diverging promoters

The applied force directed sliding by post-termination complexes and determined at what promoter re-initiation occurred. Consequently, post-termination complexes slid back to re-initiate multiple times at promoters oriented against the force but were directed away from a promoter oriented with the force (Fig. 3a-d). In a template with convergent promoters, the promoter aligned with the force was utilized only if an RNAP located this promoter at the initial round (Fig. 3a&b). In a template with divergent promoters, the promoter aligned with the force was utilized either during the initial round or during the last round of recycling (Fig. 3c&d). This result suggests that very slight forces affecting post-termination complex sliding might bias transcription from nearby but oppositely oriented promoters *in vivo*.

**Figure 3.**
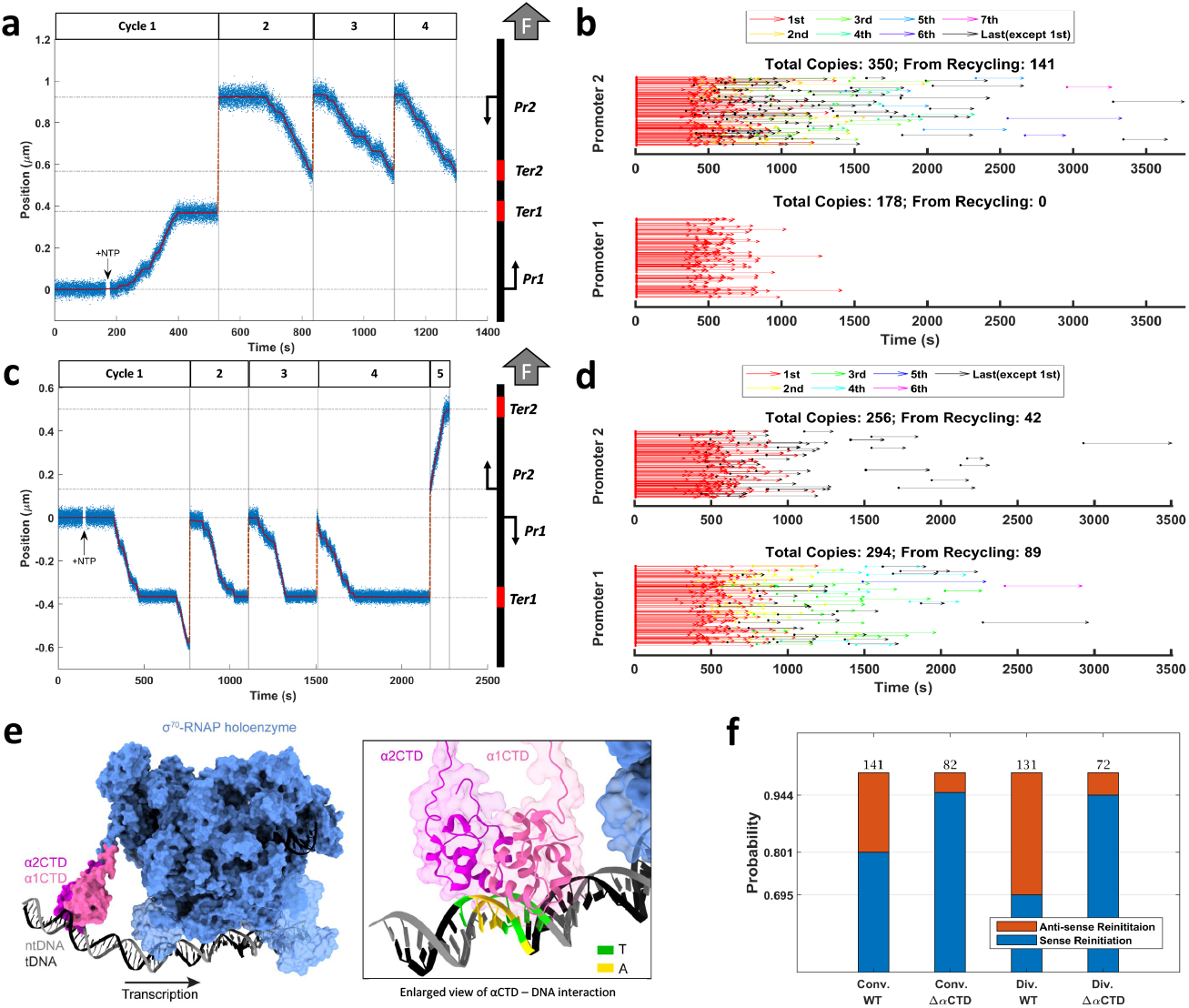
Repetitive transcription on templates with two adjacent promoters. (a) In a recording for a template with converging promoters, after one transcription event from P1 under assisting force, RNAP slid to P2 and completed three cycles of transcription opposing force before finally dissociating from the promoter. (b) A time-based catalog of the transcription events involving promoter P2 observed along the templates with converging promoters shows the beginning and end of each transcription event with different rounds depicted in different colors. Transcription from P1, the promoter oriented in the direction of force, never repeated. (c) In a recording for a template with diverging promoters, transcription from P1, opposing the direction of force, repeated four times before RNAP slid past P1 to re-initiate once from P2 assisted by force. (d) A time-based catalog of the transcription events involving promoter P2 for templates with diverging promoters shows repetitive events from promoter P1, oriented against the force but only single primary or final repetitive events from P2 oriented with the force. (e) In figures prepared with Chimera version 1.2, *α*-CTDs are shown to interact with promoter DNA in a closed *rrnB* promoter complex [18] (left). An enlarged view of the *α*-CTD shows contacts with the UP element (right). (f) Deletion of *α*-CTD diminished the probability that RNAP turned around to re-initiate in the direction opposite to the primary transcription event (anti-sense re-initiation) on templates with converging or diverging promoters.

As reported previously [6], sliding post-termination complexes readily recognized promoters with either orientation. Remarkably, the experiments described here with convergent or divergent pairs of promoters revealed equivalent dwell times before recommencing transcription in either direction (Fig. S2d). To re-initiate secondary transcription from a promoter oriented in the opposite direction, a post-termination complex, which likely has lost all or part of the transcription bubble [19], rapidly slides in the direction of applied force until it recognizes a promoter and switches polarity, seizing as the template what was previously the non-template DNA strand to form an open bubble. During this switch, nonspecific DNA contacts made by core RNAP and *σ* factor must be transiently lost. How does the enzyme remain associated with the DNA while making a U-turn? The C-terminal domains of the *α* subunits (*α*-CTD) could mediate the polarity switch. The *α*-CTDs are connected to the *α* N-terminal domains (NTDs) via long flexible linkers; the NTDs interact with the *β* (*α*1) and *β*’ (*α*2) subunits, serving as a scaffold for core enzyme assembly [20]. The *α*-CTDs make direct contacts to AT-rich sequences (UP elements) upstream of the core promoter (Fig. 3e) [18], and are required for the exceptional strength of rRNA promoters, but are dispensable for initiation at many promoters [21]. A “consensus” UP element is composed of T- and A-tracks centered at the −50 and −40 promoter positions [21]. T-tracks are also key signature motifs of intrinsic terminators, which may also contain a matching A-track upstream of a hairpin [22]. Thus, the *α*-CTDs may maintain contacts with their preferred DNA elements in the course of polarity switch, anchoring RNAP on the DNA despite the loss of other protein-DNA contacts. To explore this possibility, we measured the probability of transcription from divergent or convergent secondary promoters using an RNAP variant lacking the *α*-CTD. This mutant RNAP rarely turned around to re-initiate transcription at promoters oriented in the direction opposite to the preceding transcription event (Fig. 3f, S3c&d). However, the missing *α*-CTD did not affect RNAP’s ability of re-initiating transcription from a promoter aligned with preceding transcription event (Table S1 convergent and divergent promoter records summary).

## 3 Discussion

These experiments shed light on a mechanism of repetitive transcription that may contribute significantly to gene regulation. A four-step mechanism from the completion of elongation of an RNA transcript to the start of a second round of transcription consists of (1) RNAP remaining associated with the DNA template after elongation, (2) RNAP sliding or diffusing along the DNA, (3) RNAP stopping at obstacles or promoters, and (4) RNAP re-initiating transcription at these sites. In step (1), the RNAP-DNA complex likely undergoes some conformational change that allows the RNAP to slide freely along DNA. This may resemble a final stage of transcription termination in which the DNA strands of the transcription bubble re-hybridize [19]. In step (2), RNAP either diffuse along the DNA as previously reported [5, 6], or slide rapidly in one direction driven by as little as sub-piconewton levels of force. In step (3), RNAP (i) recognize promoters in either orientation or only aligned in the direction of the previous elongation if the *α*-CTDs are missing or (ii) become blocked by DNA-bound proteins. In step (4), RNAP can form open complexes and initiate transcription at promoters with a characteristic delay of about a minute, or even at a non-promoter site if roadblocked for 5-fold longer. The presence of DNA-bound proteins, *σ* factors and tension would significantly bias the products of the secondary transcription, as illustrated in Fig. 4.

**Figure 4.**
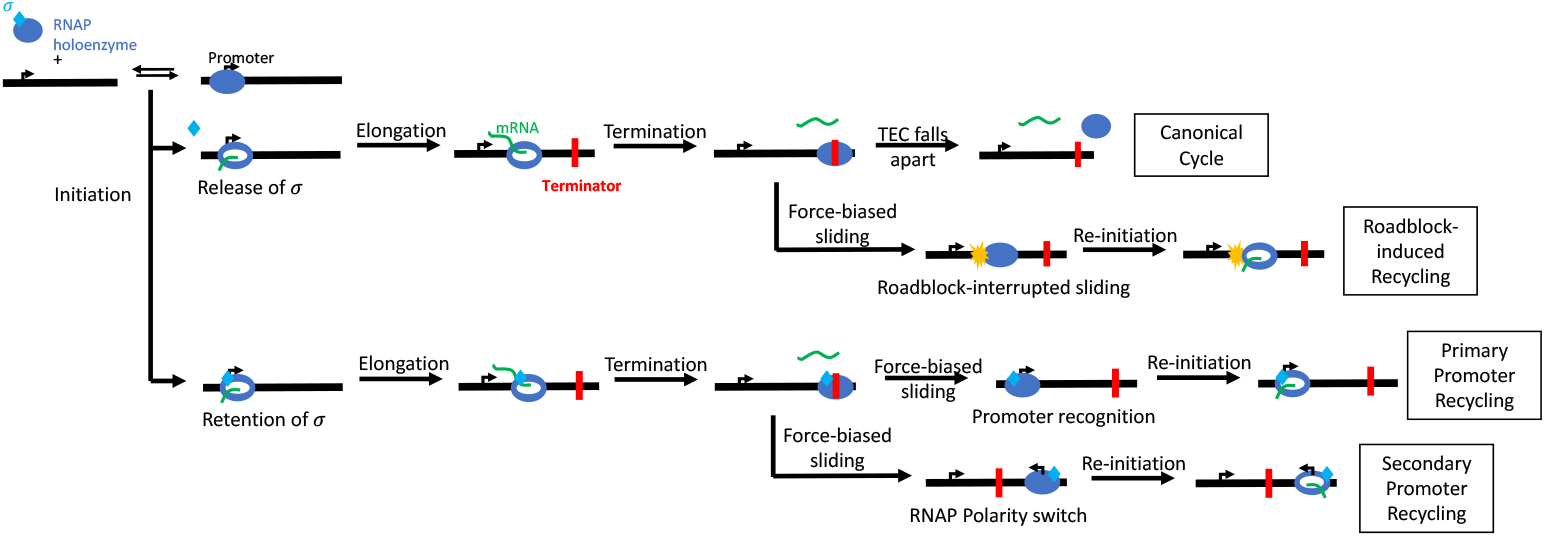
The post-termination fate of RNAP includes secondary transcription. After termination, RNAP (blue) may release mRNA (green) and dissociate from the DNA, or slide along the template. If blocked by a protein roadblock (yellow), RNAP may re-open a transcription bubble at the roadblock site and exhibit roadblock-induced recycling, a process independent of the presence of *σ* factor (cyan). Upon recognition of a distant promoter with the help of *σ* factor, wild-type RNAP may re-initiate transcription at promoters oriented in either direction.

Subtle force opposing or assisting RNAP translocation and DNA-bound proteins (roadblocks) along the template could greatly affect the efficiency and output of repetitive transcription. Even a tiny amount of force greatly reduces the time required to slide to a re-initiation site. Moreover, force restricts sliding to a single direction along a DNA sequence. Opposing force increases the frequency of repetitive transcription of a single promoter by an individual RNAP and necessarily diminishes gene expression from other promoters further downstream. Assisting force prohibits repetitive transcription and drives a sliding post-termination complex to a downstream promoter. Thus the direction of force acting on RNAP might modulate the expression of one or more genes. Indeed, it biases transcription between convergent promoters without transcriptional interference between RNAPs initiating from opposing promoters [23, 24]. The collision model would not predict similar levels of interference between divergent genes. In contrast, force-directed RNAP diffusion produces preferential expression of one transcript independently of the arrangement of promoters or additional regulatory factors [25]. Even factors like RapA, which was recently shown to accelerate dissociation of post-termination complexes [10] and would therefore diminish post-elongation sliding, would not weaken preferential, force-driven expression.

In absence of free *σ* factor in solution, less than 10% of post-termination complexes exhibited repetitive transcription at a promoter site, which is likely the portion of post-termination complexes retaining *σ* factor [9]. Adding NusG reduced that ratio to *∼*2% suggesting that retention of *σ* factor enhances sliding and repetitive transcription. In the absence of force, high affinity roadblocks would limit the sequences accessible to diffusing RNAPs and increase dwell times. However, a sliding RNAP driven by force against a high affinity roadblock might pause long enough to start transcription with or without *σ* factor. This mechanism may differ from canonical transcription initiation in which the *σ* factor assists promoter recognition and DNA strand separation. How a core RNAP enzyme might form an open bubble remains to be investigated.

It is curious that post-termination sliding has not been previously reported in force spectroscopy assays of transcription [12, 13, 17, 26, 27] in which post-termination complexes mostly dissociated from the template. One might hypothesize that in these experiments, RNAP slid too fast to seize the promoter and ran off the end of the template. However, the current experiments show that post-termination complexes often pause for several seconds at the terminator before sliding at as much as 500 bp/sec driven by forces as high as 5 pN. At this sliding rate they also readily seize promoters and re-initiate transcription. This finding contradicts the previous assumption that 3 pN of tension could make sliding too fast and transient to be detected [5]. Alternatively, higher forces employed in much of the previous work might disrupt weak interactions between a sliding RNAP and the DNA backbone, although optically resolved sliding was not observed in transcriptional interference assays without force either [28]. Most previous measurements utilized optical trapping with a feedback mechanism to maintain constant force. This may be suitable to monitor slower processive steps of molecular motors (transcribing RNAPs), but high bandwidth feedback may be critical for faster, continuous, non-processive events (sliding RNAPs) [29].

Force significantly directed sliding of post-termination complexes to accelerate repetitive transcription and modulate the relative utilization of adjacent promoters. Stringent control of the constituents afforded by the single molecule assembly revealed that *σ*^70^ often remained associated with the RNAP core enzyme to enhance sliding. Furthermore, DNA-binding proteins acting as roadblocks to sliding established non-promoter locations at which RNAP re-initiated transcription with five-fold greater delays than those observed for promoters. Sliding post-termination complexes indiscriminately utilized promoters oriented in either direction but could not easily switch template strands once the *α*-CTDs were deleted. These experiments highlight how very slight forces affecting post-termination diffusion of RNAP significantly impact transcription regulation.

## 4 Materials and Methods

### 4.1 *E. coli* Biotinylated RNAP Holoenzyme

Plasmids encoding wild-type *E. coli* core RNAP [pIA1202; *α*-*β*-*β*’[AVI][His]-*ω*) or Δ-*α*CTD RNAP [pIA1558; *α*1-235-*β*-*β*’[AVI][His]-*ω*) under control of the phage T7 promoter were transformed into *E. coli* BL21(*λ*DE3) and grown in LB at 37°C to OD600 *∼*0.5. Protein expression was induced with 0.5 mM IPTG for 5 hours at 30°C. To increase the efficiency of RNAP biotinylation, we separately expressed *E. coli* biotin ligase BirA from the T7 promoter (Addgene#109424) in E. coli BL21(*λ*DE3) using the same induction conditions. Cells were pelleted by centrifugation (6000 x g, 4°C, 10 min). Cell pellets containing overexpressed RNAP and BirA were mixed and resuspended in Lysis Buffer (10 mM Tris-OAc pH 7.8, 0.1 M NaCl, 10 mM ATP, 10 mM MgOAc, 100 µM d-biotin, 5 mM *β*-ME) supplied with Complete EDTA-free Protease Inhibitors (Roche) per manufacturer’s instructions.

Cells were open by sonication. Cell debris was pelleted by centrifugation (20,000 x g, 40 min, 4°C). The cleared cell extract was incubated with Ni Sepharose 6 Fast Flow resin (Cytiva, Marlborough, MA) for 40 min at 4°C with agitation. The resin was washed with Ni-A Buffer (25 mM Tris pH 6.9, 5% glycerol, 500 mM NaCl, 5 mM *β*-ME, 0.1 mM phenylmethylsulfonyl fluoride (PMSF) supplemented with 10 mM, 20 mM, and 30 mM imidazole. Protein was eluted in Ni-B Buffer (25 mM Tris pH 6.9, 5% glycerol, 5 mM *β*-ME, 0.1 mM PMSF, 100 mM NaCl, 300 mM imidazole).

The sample was diluted 1.5 times with Hep-A Buffer (25 mM Tris pH 6.9, 5% glycerol, 5 mM *β*-ME) and then loaded onto Heparin HP column (Cytiva). A linear gradient between Hep-A and Hep-B Buffer (25 mM Tris pH 6.9, 5% glycerol, 5 mM *β*-ME, 1 M NaCl) was applied. The biotinylated RNAP core was eluted at *∼*40 mS/cm.

The elute from Heparin HP column was diluted 2.5 times with Hep-A Buffer and loaded onto Resource Q column (Cytiva). A linear gradient was applied from 5% - 100% Hep-B Buffer. The biotinylated RNAP core was eluted at *∼*25 mS/cm.

Fractions from the elution peaks were analyzed by SDS-PAGE. Those containing purified protein were combined and dialyzed against Storage Buffer (20 mM Tris-HCl, pH 7.5, 150 mM NaCl, 45% glycerol, 5 mM *β*-ME, 0.2 mM EDTA).

*E. coli* RNAP holoenzymes were reconstituted from *σ*^70^ initiation factor, expressed as previously described [30], and the biotinylated RNAP core enzyme.

### 4.2 Lac Repressor Protein

Lac repressor protein (LacI) was prepared in the laboratory of Kathleen Matthews as previously described [31].

### 4.3 Transcription templates

The DNA templates for single promoter opposing and assisting force transcription assays were PCR amplicons from a plasmid template containing the T7A1 promoter followed by 23 bp without G in the template strand, a 1.2 kb downstream elongation region including a lac operator site (O1), and finally a terminator sequence (*λ*T1, t500, or trpA1-*λ*T1; Supplementary Information). The plasmid template was amplified with single-digoxigenin labeled forward and unlabeled reverse primer pairs, and Q5 Hot Start High-Fidelity 2X PCR Master Mix (New England Biolabs, Ipswich, MA). The transcribed region had the following spacings: Promoter-713 bp-Lac operator-612 bp-Terminator, as illustrated in Figure 1b. For the opposing force experiments, primers D-A/JBOIDO1/5096 and S/JBOIDO1/2086 were used to generate a 3k bp amplicon with 1021 bp between the chamber surface anchor point and the transcription start site. For the assisting force experiments, primers D-S/YY-400-103 and A/pUC18-nuB104/2043 were used to generate a 4k bp amplicon with 2014 bp between the anchor point and transcription start site. The longer separation in the assisting force amplicon reduces adhesion of promoter-bound magnetic beads to the chamber surface at the beginning of transcription.

In preparation of the templates with the convergent promoters, we used a digoxigenin-labeled primer D-S/YY-400-103 and a primer with the palindromic restriction site ApaI A/pUC18-nuB104/1983-ApaI to generate amplicons with digoxigenin label on one end and ApaI restriction site on the other end. We repeated the PCR with unlabeled primer S/YY-400-103 and the same ApaI containing primer to generate amplicons with the same restriction site but no digoxigenin label. Both amplicons were then digested by ApaI enzymes (New England BioLabs) in rCutSmart buffer (New England Labs) per manufacturer’s instruction. To produce satisfactory amount of constructs with a digoxigenin-labeled end and an unlabeled end, the digoxigenin-labeled and unlabeled amplicons were ligated with T7 ligase (New England BioLabs) in a specific ratio of 1:2. Similary, in preparation of the templates with the divergent promoters, we paired a digoxigenin-labeled primer D-A/JBOIDO1/5096 and its unlabeled version with a primer with palindromic SalI restriction site S/JBOIDO1/2086-SalI to generate two sets of amplicons, which were then digested by SalI (New England Labs) restriction enzymes in NEBuffer r3.1 (New England Labs) and ligated in 1:2 ratio to generate the divergent promoter templates. The sequences of all plasmids and primers used in these experiments are listed in Supplementary Information (TableS2).

### 4.4 Preparation of halted TEC and tethers

Stalled complexes were prepared by incubating 1.5 nM DNA with 15 nM RNAP in Transcription Buffer (20 mM Tris glutamate pH = 8, 50 mM potassium glutamate, 10 mM magnesium glutamate, 1 mM DTT, 0.2 mg/ml *α*-casein) for 20 min at 37°C. After introducing 0.5 mM of ATP, UTP and GTP, RNAPs transcribed 23 bases and then stalled at the first *G* in the template.

Microchambers were incubated with 8 mg/µl anti-digoxigenin (Roche) at 25°C for 90 min and then the blocking buffer (PBS with 1% *α*-casein) at 25°C for 30 min. Then, halted complexes were introduced into the anti-digoxigenin-coated microchamber, incubated at 25°C for 20 min to form tethers. Streptavidin-coated paramagnetic beads (Dynbeads MyOneT1, Thermofisher, Waltham, MA) were introduced last to the chamber and incubated with tethers at 25°C for 3 min. Chambers were washed with excessive (4 volumes) Transcription Buffer to remove free RNAP molecules and beads in solution before recording.

### 4.5 Magnetic Tweezers (MT) Assays

Transcription complexes were prepared with E. coli RNAP holoenzyme biotinylated on the C-terminus of the *β*’ subunit and DNA molecules labeled at one end with digoxigenin for anchoring to the microchamber surface. These molecules included a T7A1 promoter, a binding site for lac repressor, and a terminator in a 1.2 kbp sequence. Each RNAP enzyme in a magnetic tweezing assembly was coupled to a streptavidin-coated magnetic bead and transcription elongation complexes were stalled after 23 bp by supplying only ATP, GTP, and UTP before tethering to the surface of an anti-digoxigenin-coated microchamber. Then the flow chamber was placed on the MT microscope stage, and a field of view with several freely moving tethered beads was selected. After a brief period of tracking the selected beads to establish a baseline tether length, 1 mM NTPs including CTP were introduced to produce transcription that increased or decreased tether lengths for the assisting, or opposing, force configurations respectively (Fig. 1a). Since the magnetite content of individual beads varies slightly and DNA tethers become extended to differing degrees, the transcription (tether length) records were scaled to assemble a data set for statistics (see Fig. S1 and Data Processing in SI).

## Acknowledgments

LacI was a generous gift from Kathleen Matthews, Rice University. Plasmids for these experiments were created by Derrica McCalla. We thank Wenxuan Xu and Yan Yan for preliminary measurements. This work was supported by the National Institutes of Health (NIH) grants R01 GM084070 and R35GM149296 to LF and R01 GM067153 to IA.

## Supplementary Information

**Figure S1.**
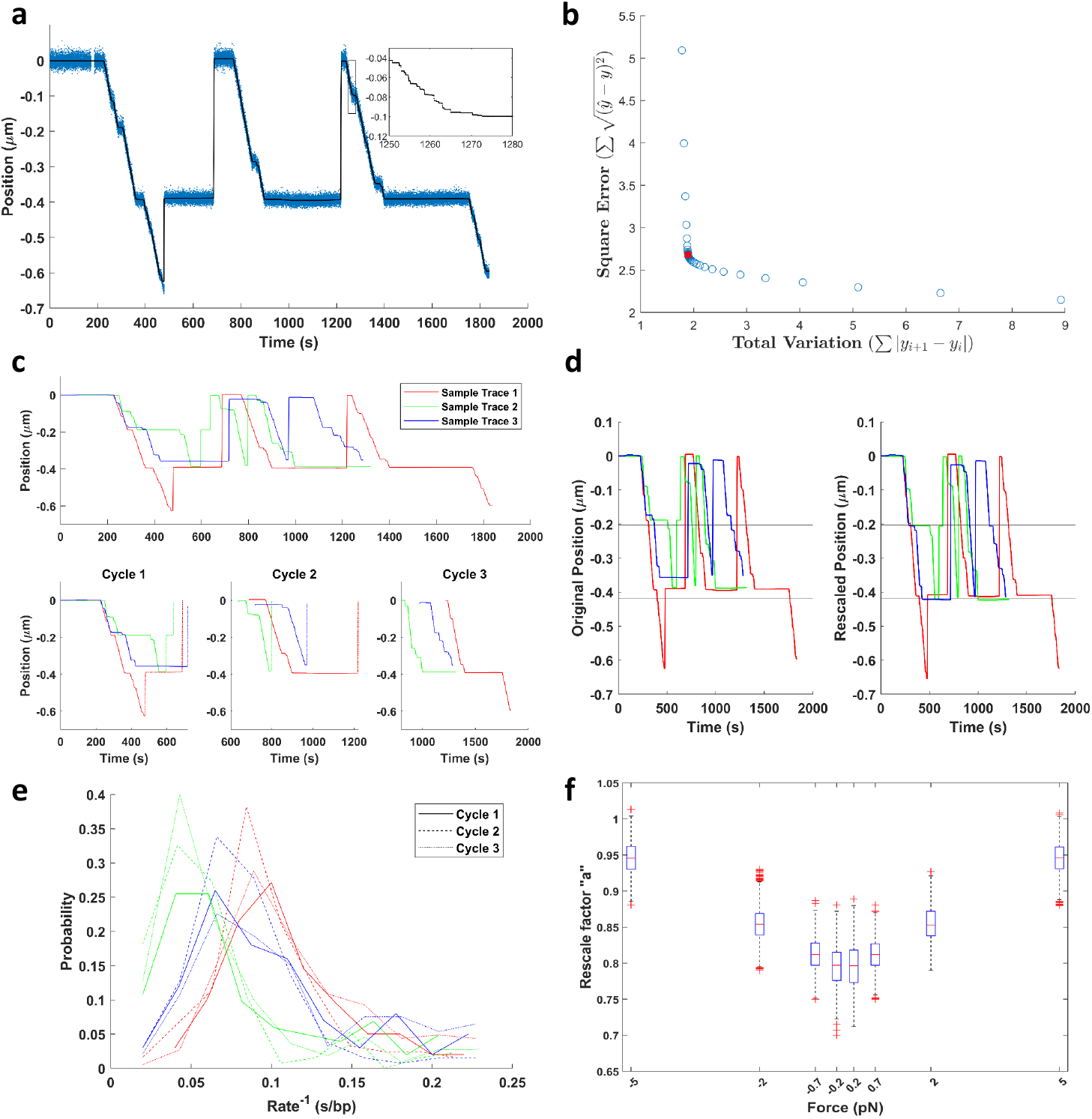
(a) A sample record with three transcription cycles (blue dots) and the de-noised result (black line). The inset shows the enlarged view of the step-wise curve after de-noising. (b) The selection of *λ* value in the de-noising process. The blue circle shows the magnitudes of the square error and the total variation term at different *λ*, and the red asterisk is the optimal *λ* value used for de-noising the trace, where two terms are balanced. (c) Three de-noised records with multiple transcription cycles plotted in different colors are shown in the upper panel. The separated cycles are shown in the lower panels. The detected sliding events are indicated by dotted lines. An entire cycle is defined as starting from RNAP promoter binding and ending after post-termination sliding. (d) Rescaled records in (c) diplay significant pauses at the operator and terminator sites. (e) The transcription rates of individual cycles from the records in (d) are plotted in different line styles and colors. The rate distributions show similar peaks within the same records but peaks differ between different record, suggesting that the multiple cycles in a single record were likely produced by a single RNAP. (f) The force vs. rescaling factor for the entire data set. As expected, rescaling factors are smaller for records under lower forces.

**Figure S2.**
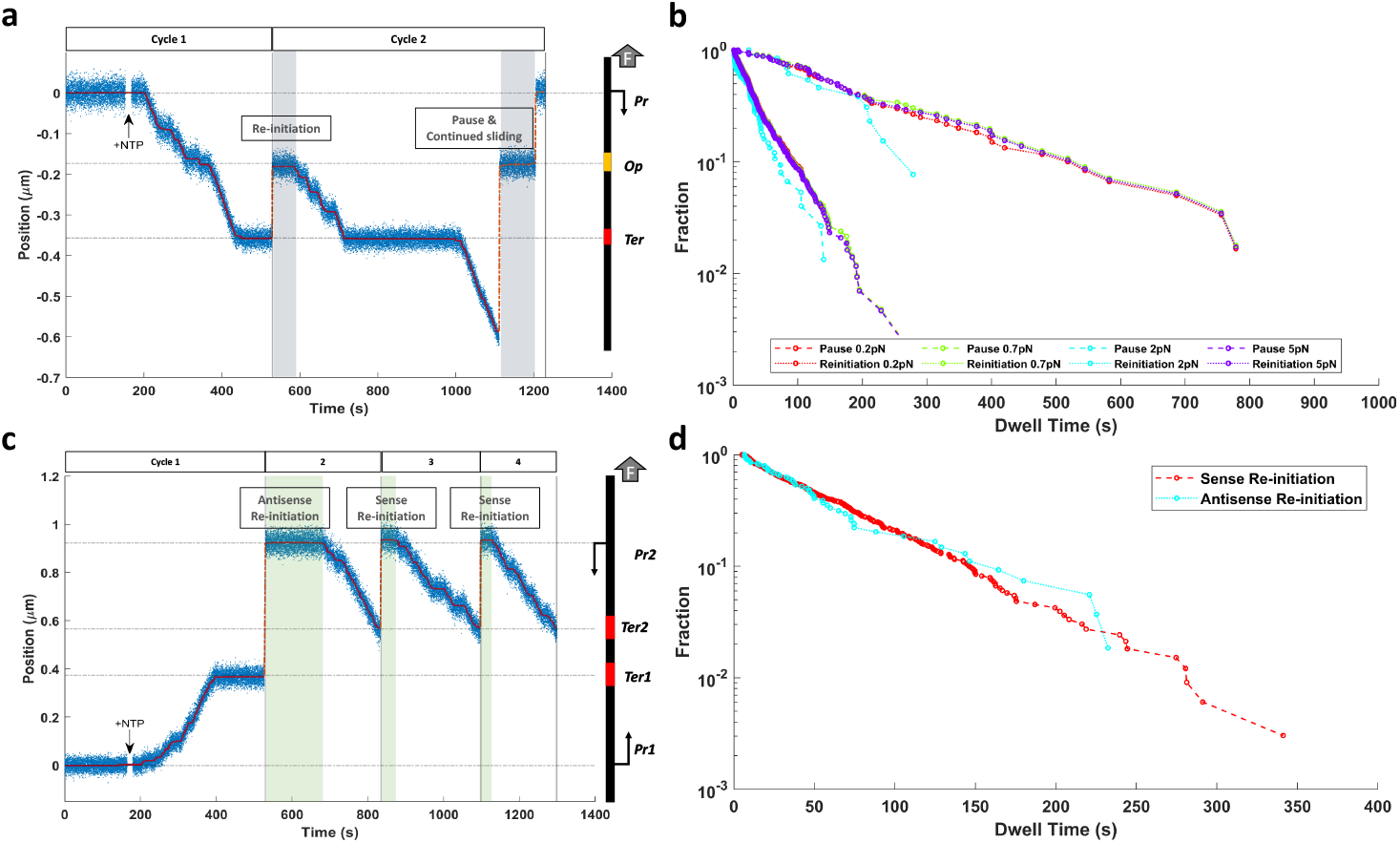
(a) A sample record shows pauses by a sliding RNAP roadblocked at the lac repressor binding site followed by re-initiation (at *∼*530 s) or continued sliding after pause (at *∼*1130 s). The shaded intervals are examples of dwell times for each of these events. (b) Dwell times at the lac repressor binding site prior to re-initiation (dotted lines) were longer and produced a much broader distribution than dwell times preceding continued sliding events (dashed lines). Remarkably, the magnitude of force had a negligible effect on these distributions. (c) A sample record shows both anti-sense and sense re-initiation events. The shaded intervals indicate dwell times at the promoter prior to re-initiation. (d) The distributions of dwell times at the promoter site were similar preceding anti-sense and sense re-initiation.

**Figure S3.**
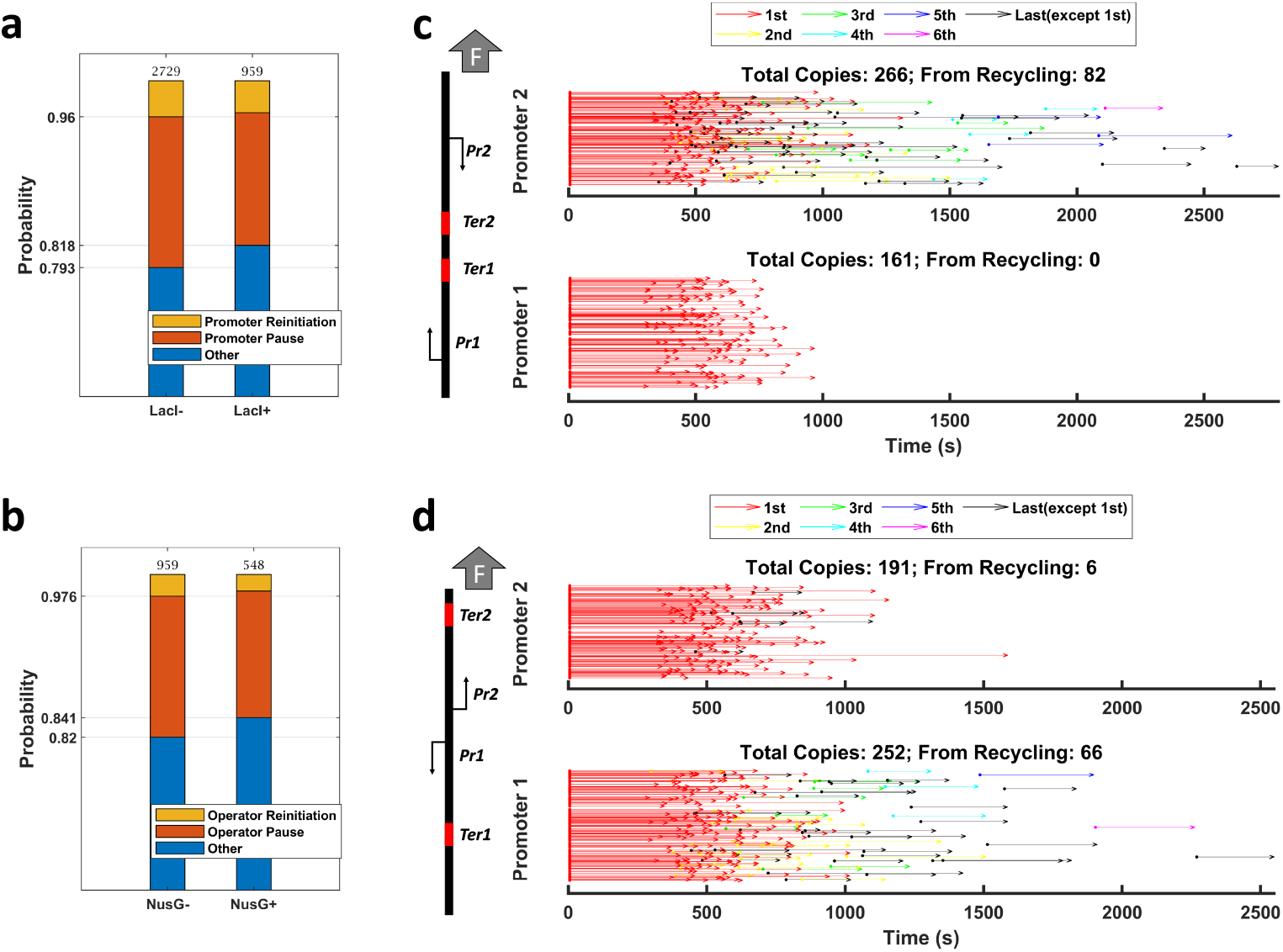
(a) The presence of LacI has a negligible effect on the probability of promoter pause and subsequent re-initiation. (b) The presence of NusG has a negligible effect on the probability of roadblocking at the lac repressor site or subsequent re-initiation. (c) A summary of Δ*α*-CTD shows RNAP transcription events on templates with convergent or (d) divergent promoters. Compared to wild-type RNAP Fig. 3b and d, the deletion of *α*-CTD reduced the ability of RNAP to turn around and re-initiate from a secondary promoter oriented in the the direction opposite to the preceding transcription event.

### Data processing

In the first step of the analysis, the initial and final sections of a record, before the addition of NTPs and after beads dissociation, were cropped out to generate clean and informative traces (Fig. 1b, 2a, 3a&c, and S1a). The zero point was set to the average of values within the first 3 s of a record.

Since the records often consist of multiple rounds of transcription cycle with sliding returns in between, the simple moving mean/median smoothing algorithm is not suitable. We used a total variation de-noising algorithm[32, 33] which relies on minimizing a two-term cost function

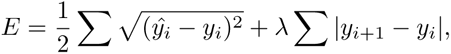

where the first term describes the square errors between the de-noised signal *y*^ and original signal *y*, and the second term describes the total variation of the de-noised signal. At different values of regularization parameters *λ*, the values of two terms were plotted in Fig. S1b. The *λ* at the turning point was then selected to produce optimal values of both terms in the cost function. At the optimized *λ* values, the total variation term is minimized to remove noise and generate monotonically increasing or decreasing signals within individual transcription and post-termination sliding events. The square error term, on the other hand, is minimized to preserve the pattern of RNAP pause and progression during elongation and post-termination sliding (Fig. S1a). The de-noised records were re-sampled to a uniform 50 Hz frequency. This re-sampling provided convenient and standardized time series for the generation of histograms in subsequent analyses.

In the next step, a contour length representation of the template was re-scaled to fit spacings between the promoter and the terminator in the de-noised data. This normalization was necessary because DNA templates attached to different beads were under slightly different tensions and therefore stretched to different extents. The template representation was scaled by a factor ‘a’ and shifted by a factor ‘b’, *y_contour_* = (1*/a*) *∗ y_exp_* + *b*, in which *y_contour_* is the expected RNAP position after rescaling according to the template contour length, and *y_exp_* is the experimentally observed RNAP position before rescaling. Since the zero position was set to the average of mean values of the first 3 seconds of a record (approximately promoter site), the factor ‘b’ was effectively zero. The factor ‘a’ was determined to be the value at which the rescaled data generated the maximum dwell time peaks at the expected positions of operator and terminator sites. After normalization, processive transcription on all records started from the promoter site (0 bp) and either terminated or exhibited a long pause at the terminator site (Fig. S1d). Figure S1f shows the symmetric distribution of the rescaling factor ‘a’ from positive to negative forces, which ranged from 0.95 at 5 pN force to 0.8 at 0.2 pN.

A record with multiple rounds of transcription must be sectioned for further analysis of individual cycles. However, sectioning is not trivial due to the step-wise and pause-interspersed nature of the de-noised curve. To reliably separate transcription events from post-termination sliding, we first forced a signal, even with post-termination sliding in the opposite direction, to be monotonically decreasing by reversing any positive changes in position. This treatment transformed the signal into a consistent downward trajectory and eliminated the need to account for the possibility of the RNAP molecule revisiting positions due to recycling. Next, we computed histogram counts with a 0.003 *µm* bin size from the monotonically decreasing signal, and assigned the most likely states from “paused”, “transcription”, “sliding” and “unknown” to each bin according to the dwell times within the bin. Then, we merged consecutive states of the same type. In particular, the “paused” and “unknown” states can mix into consecutive “transcription” or “sliding” states. After this step, the signal was separated into multiple sections, with “transcription” and “sliding” states alternating. Finally, we removed states with insignificant movement (*<*0.05 *µm*) or fleeting duration (“transcription” state *<*10 s) from the sections and adjusted the start and end times of the neighboring sections if necessary to ensure a seamless flow between states. Figure S1c shows that three sample traces, each spanning three rounds of transcription cycles, were faithfully separated, and the post-termination sliding (dotted lines) was clearly differentiated from transcription (solid line). This protocol accurately sectioned all records regardless of the orientations of transcription and post-termination sliding.

### Analysis of processed data

The processed time series consisted of alternating “transcription” and “sliding” sections. Within these sections, the time series were always monotonically increasing/decreasing, and the paused sites were exactly flat. This systematic structure enabled the transformation of each segment into a histogram. From these histograms, we can extract pivotal information such as pause sites, durations, and start and end positions of post-termination sliding. In particular, the recycling ratios at different terminators and force conditions (Fig. 1c & d) denoted the proportion of records where a sliding section succeeded the initial transcriptional phase. The RNAP post-termination sliding rates (Fig. 1e) were determined by measuring the duration spent in sliding over the total distance covered, excluding pauses exceeding 1 second. The start and end times/positions of a sliding section were extracted from the start and end points of a sliding event. Notably, the presence of pauses within a sliding section bifurcated a sliding event into distinct segments. The first segment concluded at the pause site, pausing momentarily before the second segment commenced from the same pause site (Fig. 2b, 3b & d, S3c & d).

In the identification of pauses induced by specific sequences or DNA-bound obstacles at promoter, operator, and terminator sites, we pinpointed the longest pauses within anticipated positions, a range defined as *±* 0.02*µm*. This criterion was justified by the inherent expectation that pauses induced by these sequence-specific or roadblock-related factors would exhibit remarkably longer characteristic dwell times in comparison to the ubiquitous pauses encountered throughout the dataset. Even in rare cases that this selection might mistakenly assign a ubiquitous pause as the sequence/roadblock-induced pause, the consequence was merely the substitution of a short sequence or roadblock-induced pause with an equally brief ubiquitous pause and thus not significantly skew the overall statistical analyses. The identified sequence/roadblock-induced pauses were used to generate Figure S2b & c.

**Table S1.**
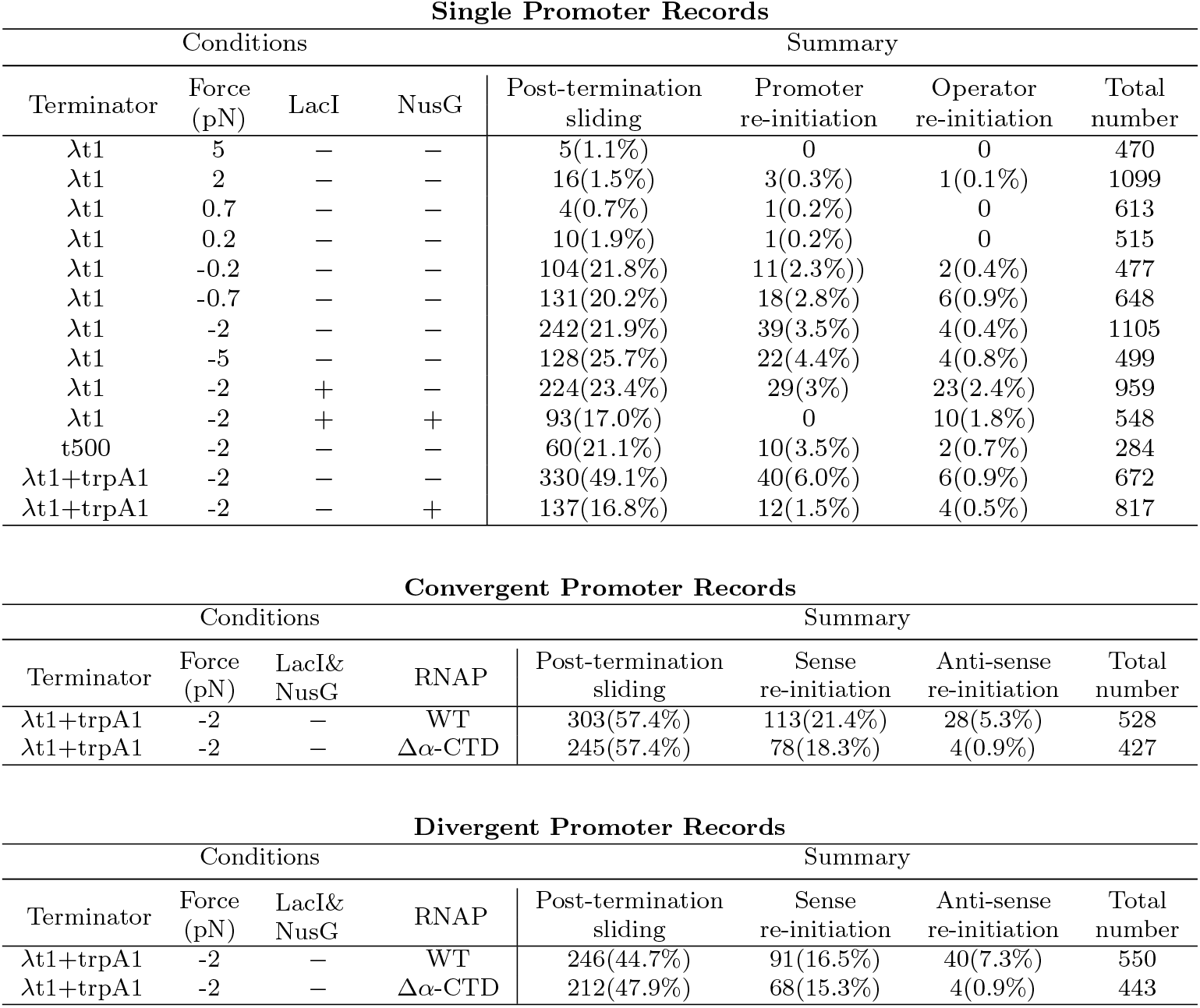
Summary of records using single promoter, convergent promoter, and divergent promoter templates. Each record was analyzed based on the post-termination fate of RNAP. Records that exhibited post-termination sliding and subsequent re-initiation of transcription were included in both the post-termination sliding and re-initiation categories.

**Table S2.**
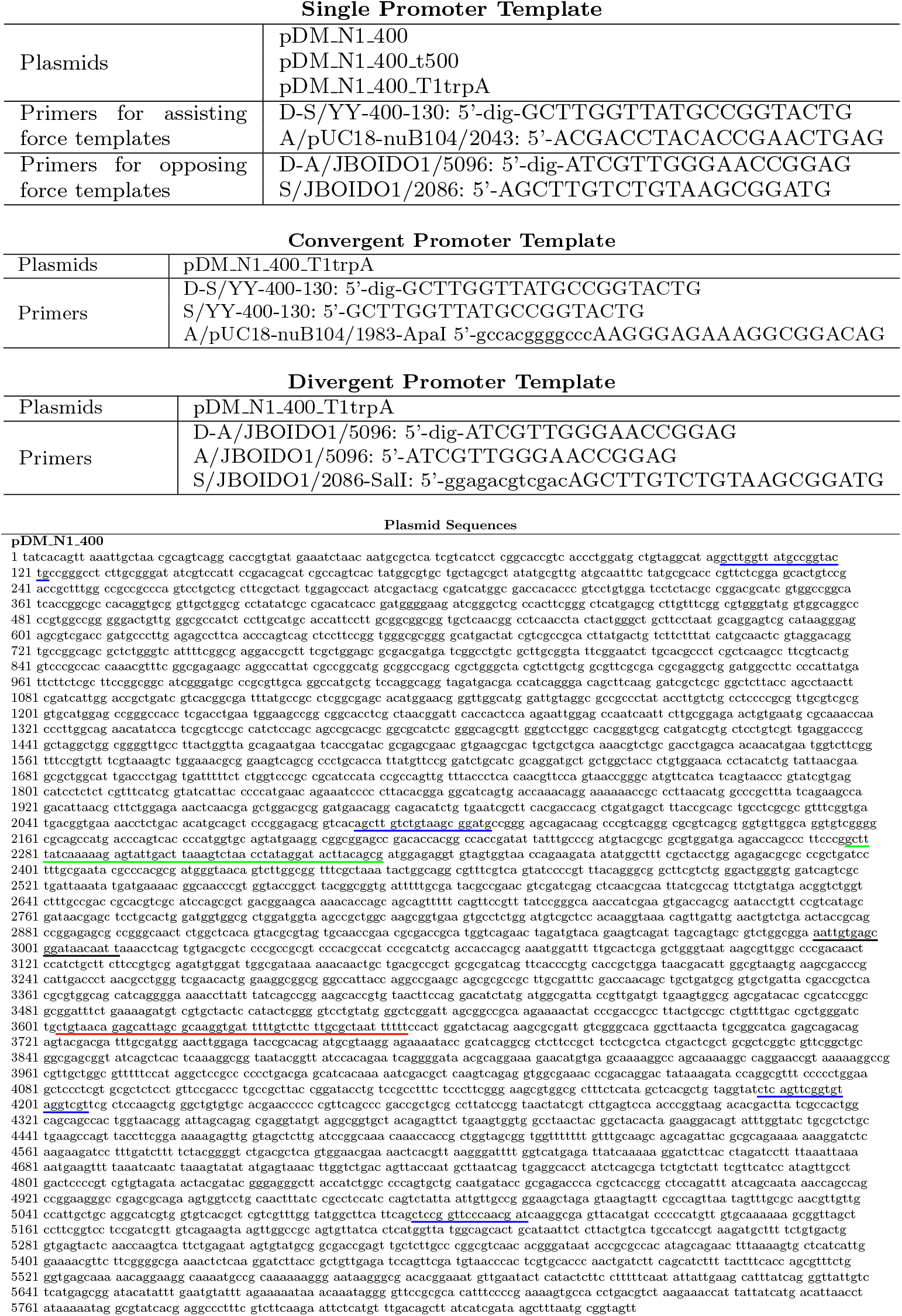
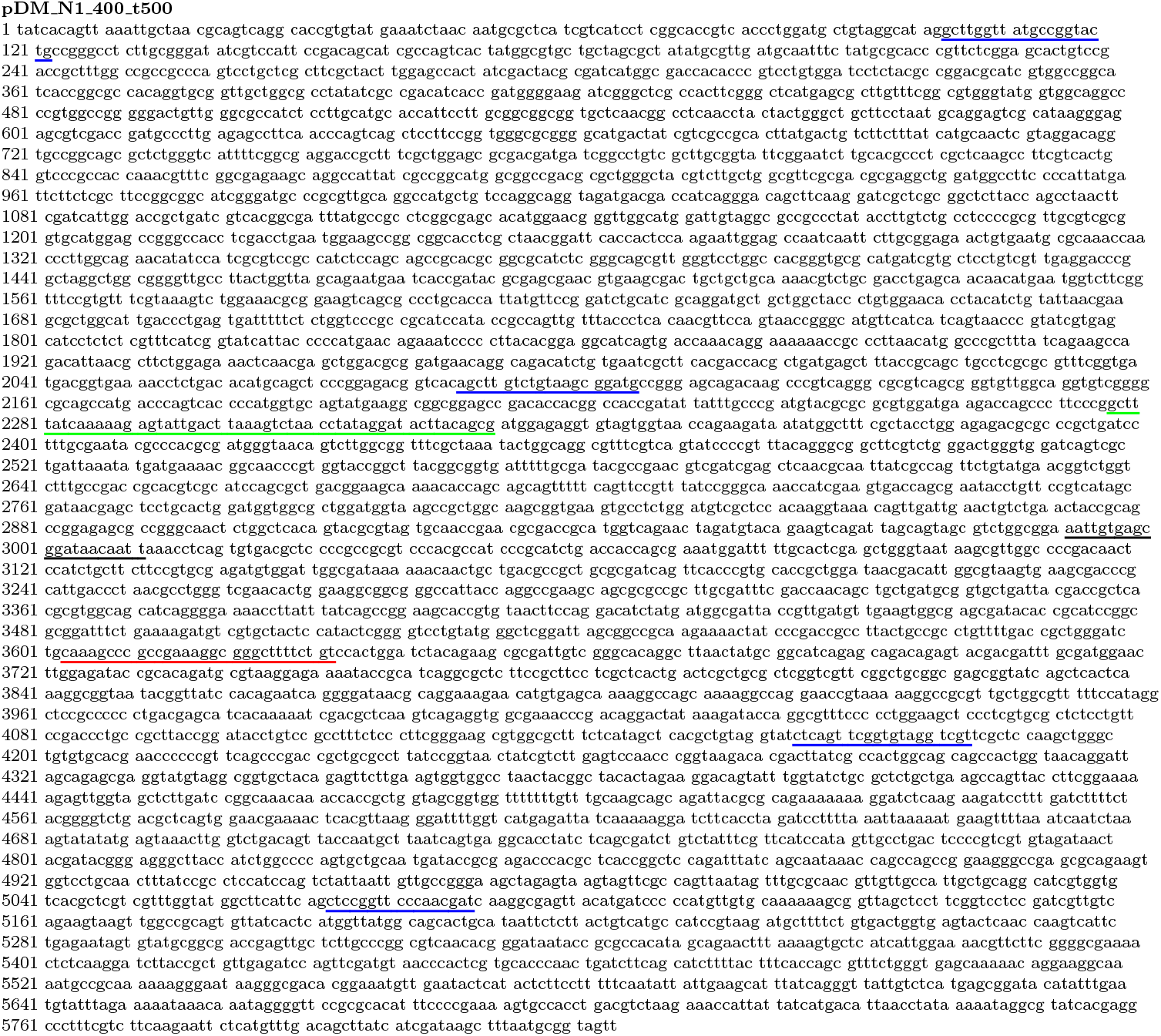
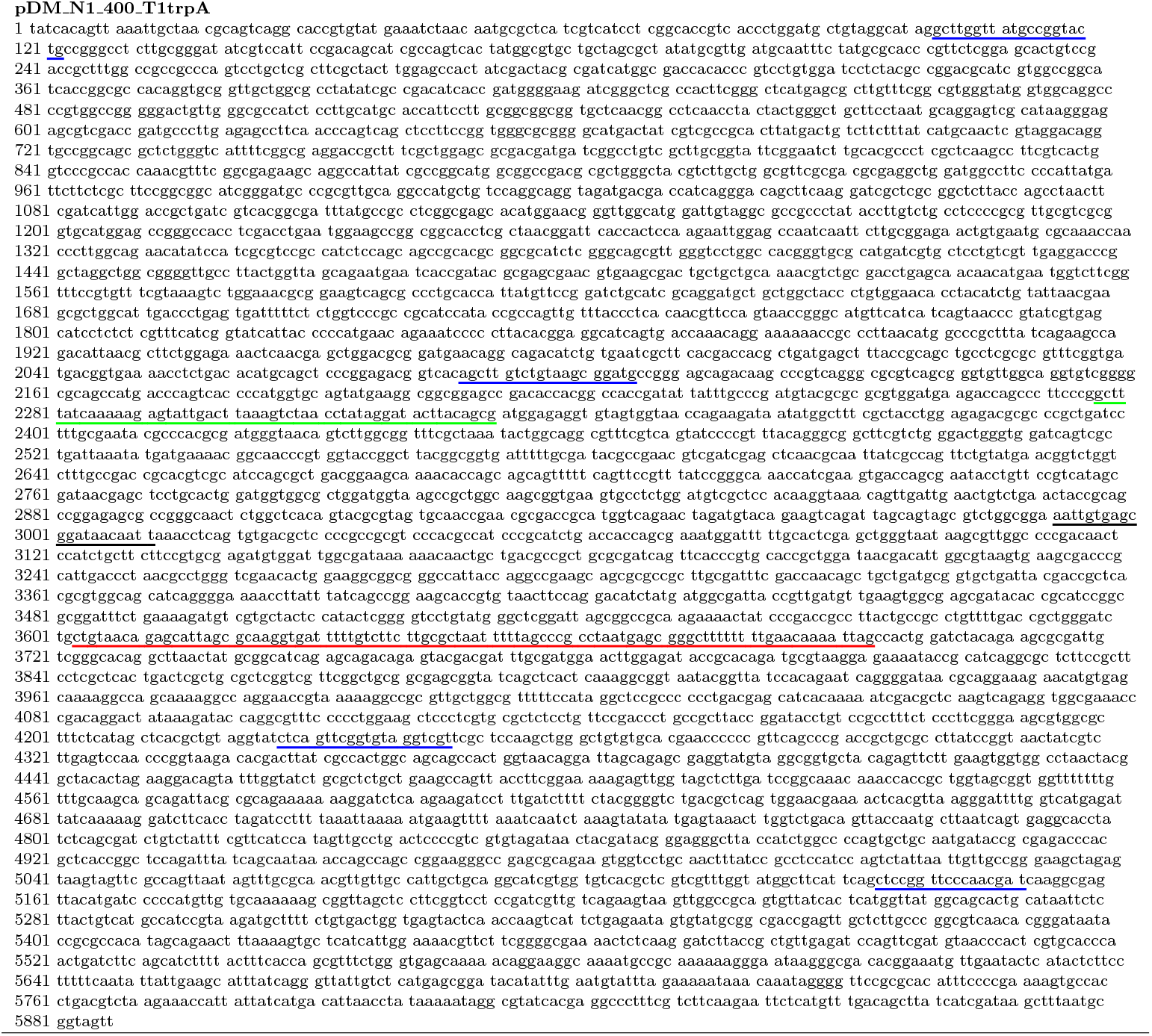
Summary of plasmids and primers used in the experiments. Colored underline: primers (blue), promoter (green), operator (black), terminator (red).

## Notes

### Competing Interest Statement

The authors have declared no competing interest.

## References

[1] Berg, O.G., Winter, R.B., Von Hippel, P.H.: Diffusion-driven mechanisms of protein translocation on nucleic acids. 1. models and theory. Biochemistry 20(24), 6929–6948 (1981)

[2] Wang, F., Redding, S., Finkelstein, I.J., Gorman, J., Reichman, D.R., Greene, E.C.: The promoter-search mechanism of escherichia coli rna polymerase is dominated by three-dimensional diffusion. Nature structural & molecular biology 20(2), 174–181 (2013)

[3] Arndt, K.M., Chamberlin, M.J.: Transcription termination in escherichia coli: measurement of the rate of enzyme release from rho-independent terminators. Journal of molecular biology 202(2), 271–285 (1988)

[4] Bellecourt, M.J., Ray-Soni, A., Harwig, A., Mooney, R.A., Landick, R.: Rna polymerase clamp movement aids dissociation from dna but is not required for rna release at intrinsic terminators. Journal of molecular biology 431(4), 696–713 (2019)

[5] Harden, T.T., Herlambang, K.S., Chamberlain, M., Lalanne, J.-B., Wells, C.D., Li, G.-W., Landick, R., Hochschild, A., Kondev, J., Gelles, J.: Alternative transcription cycle for bacterial rna polymerase. Nature Communications 11(1), 448 (2020)

[6] Kang, W., Hwang, S., Kang, J.Y., Kang, C., Hohng, S.: Hopping and flipping of rna polymerase on dna during recycling for reinitiation after intrinsic termination in bacterial transcription. International Journal of Molecular Sciences 22(5), 2398 (2021)

[7] Kang, W., Ha, K.S., Uhm, H., Park, K., Lee, J.Y., Hohng, S., Kang, C.: Transcription reinitiation by recycling rna polymerase that diffuses on dna after releasing terminated rna. Nature Communications 11(1), 450 (2020)

[8] Paget, M.S., Helmann, J.D.: The *σ*70family of sigma factors. Genome biology 4(1), 1–6 (2003)

[9] Harden, T.T., Wells, C.D., Friedman, L.J., Landick, R., Hochschild, A., Kondev, J., Gelles, J.: Bacterial rna polymerase can retain *σ*70 throughout transcription. Proceedings of the National Academy of Sciences 113(3), 602–607 (2016)

[10] Inlow, K., Tenenbaum, D., Friedman, L.J., Kondev, J., Gelles, J.: Recycling of bacterial rna polymerase by the swi2/snf2 atpase rapa. Proceedings of the National Academy of Sciences 120(28), 2303849120 (2023) 10.1073/ pnas.2303849120

[11] Herbert, K.M., Zhou, J., Mooney, R.A., Porta, A.L., Landick, R., Block, S.M.: E. coli nusg inhibits backtracking and accelerates pause-free transcription by promoting forward translocation of rna polymerase. Journal of Molecular Biology 399(1), 17–30 (2010) 10.1016/j.jmb.2010.03.051

[12] Forde, N.R., Izhaky, D., Woodcock, G.R., Wuite, G.J., Bustamante, C.: Using mechanical force to probe the mechanism of pausing and arrest during continuous elongation by escherichia coli rna polymerase. Proceedings of the National Academy of Sciences U S A 99(18), 11682–7 (2002) 10.1073/pnas. 142417799

[13] Adelman, K., La Porta, A., Santangelo, T.J., Lis, J.T., Roberts, J.W., Wang, M.D.: Single molecule analysis of rna polymerase elongation reveals uniform kinetic behavior. Proceedings of the National Academy of Sciences 99(21), 13538–13543 (2002) 10.1073/pnas.212358999

[14] Chen, L., Conlisk, A.T.: Forces affecting double-stranded dna translocation through synthetic nanopores. Biomedical Microdevices 13(2), 403–414 (2011) 10.1007/s10544-011-9509-7

[15] Qian, J., Cartee, A., Xu, W., Yan, Y., Wang, B., Artsimovitch, I., Dunlap, D., Finzi, L.: Reciprocating rna polymerase batters through roadblocks. bioRxiv, 2023–0104522798 (2023) 10.1101/2023.01.04.522798

[16] NandyMazumdar, M., Nedialkov, Y., Svetlov, D., Sevostyanova, A., Belogurov, G.A., Artsimovitch, I.: Rna polymerase gate loop guides the nontemplate dna strand in transcription complexes. Proceedings of the National Academy of Sciences of the United States of America 113(52), 14994–14999 (2016) 10.1073/pnas.1613673114

[17] Larson, M.H., Greenleaf, W.J., Landick, R., Block, S.M.: Applied force reveals 14 mechanistic and energetic details of transcription termination. Cell 132(6), 971– 982 (2008)

[18] Shin, Y., Qayyum, M.Z., Pupov, D., Esyunina, D., Kulbachinskiy, A., Murakami, K.S.: Structural basis of ribosomal rna transcription regulation. Nature Communications 12(1), 528 (2021) 10.1038/s41467-020-20776-y

[19] You, L., Omollo, E.O., Yu, C., Mooney, R.A., Shi, J., Shen, L., Wu, X., Wen, A., He, D., Zeng, Y., Feng, Y., Landick, R., Zhang, Y.: Structural basis for intrinsic transcription termination. Nature (2023) 10.1038/ s41586-022-05604-1

[20] Murayama, S., Ishikawa, S., Chumsakul, O., Ogasawara, N., Oshima, T.: The role of -ctd in the genome-wide transcriptional regulation of the bacillus subtilis cells. PLOS ONE 10(7), 0131588 (2015) 10.1371/journal.pone.0131588

[21] Gourse, R.L., Ross, W., Gaal, T.: Ups and downs in bacterial transcription initiation: the role of the alpha subunit of rna polymerase in promoter recognition. Molecular Microbiology 37(4), 687–695 (2000) 10.1046/j.1365-2958.2000.01972.x

[22] Roberts, J.W.: Mechanisms of bacterial transcription termination. Journal of Molecular Biology 431(20), 4030–4039 (2019) 10.1016/j.jmb.2019.04.003

[23] Hao, N., Palmer, A.C., Ahlgren-Berg, A., Shearwin, K.E., Dodd, I.B.: The role of repressor kinetics in relief of transcriptional interference between convergent promoters. Nucleic Acids Research 44(14), 6625–6638 (2016) 10.1093/nar/gkw600

[24] Brophy, J.A.N., Voigt, C.A.: Antisense transcription as a tool to tune gene expression. Molecular Systems Biology 12(1), 854 (2016) 10.15252/msb.20156540

[25] Wu, A.C.K., Patel, H., Chia, M., Moretto, F., Frith, D., Snijders, A.P., Werven, F.J.: Repression of divergent noncoding transcription by a sequence-specific transcription factor. Molecular Cell 72(6), 942–9547 (2018) 10.1016/j.molcel.2018.10.018

[26] Davenport, R.J., Wuite, G.J., Landick, R., Bustamante, C.: Single-molecule study of transcriptional pausing and arrest by e. coli rna polymerase. Science 287(5462), 2497–500 (2000)

[27] Shaevitz, J.W., Abbondanzieri, E.A., Landick, R., Block, S.M.: Backtracking by single rna polymerase molecules observed at near-base-pair resolution. Nature 426(6967), 684–687 (2003) 10.1038/nature02191

[28] Wang, L., Watters, J.W., Ju, X., Lu, G., Liu, S.: Head-on and co-directional rna polymerase collisions orchestrate bidirectional transcription termination. Molecular Cell 83(7), 1153–11644 (2023) 10.1016/j.molcel.2023.02.017

[29] Rice, S.E., Purcell, T.J., Spudich, J.A.: [6] building and using optical traps to study properties of molecular motors. In: Methods in Enzymology vol. 361, pp. 112–133. Elsevier,(2003)

[30] Svetlov, V., Artsimovitch, I.: Purification of bacterial rna polymerase: tools and protocols. Methods in molecular biology (Clifton, N.J.) 1276, 13–29 (2015) 10.1007/978-1-4939-2392-2_2. 25665556[pmid] Methods Mol Biol

[31] Xu, J., Liu, S., Chen, M., Ma, J., Matthews, K.S.: Altering residues n125 and d149 impacts sugar effector binding and allosteric parameters in escherichia coli lactose repressor. Biochemistry 50(42), 9002–9013 (2011) 10.1021/bi200896t. doi: 10.1021/bi200896t

32. Little, M.A., Jones, N.S.: Sparse bayesian step-filtering for high-throughput analysis of molecular machine dynamics. In: 2010 IEEE International Conference on Acoustics, Speech and Signal Processing, pp. 4162–4165 (2010). 10.1109/ICASSP.2010.5495722

[33] Qian, J., Collette, D., Finzi, L., Dunlap, D.: In: Heller, I., Dulin, D., Peterman, E.J.G. (eds.) Detecting DNA Loops Using Tethered Particle Motion, pp. 451–466. Springer, New York, NY (2024). 10.1007/978-1-0716-3377-9_21.

